# Lyophilized yeast powder for adjuvant free thermostable vaccine delivery

**DOI:** 10.1101/2020.11.30.401885

**Authors:** Ravinder Kumar, Bhushan N. Kharbikar

**Affiliations:** Department of Obstetrics, Gynecology and Reproductive Sciences, University of California San Francisco, California - 94143, USA; Department of Bioengineering and Therapeutic Sciences, University of California San Francisco, California, 94158, USA

**Keywords:** Yeast-based vaccine, *P. pastoris*, long term-stability, lyophilized, yeast powder, thermostable

## Abstract

Thermolabile nature of commercially available vaccines necessitates their storage, transportation and dissemination under refrigerated condition. Maintenance of continuous cold chain at every step increases the final cost of vaccines. Any breach in the cold chain, even for a short duration results in the need to discard the vaccine. As a result, there is a pressing need for the development of thermostable vaccines. In this proof of concept study, we showed that *E. coli* curli-GFP fusion protein remains stable in freeze-dried yeast powder for more than a 13 and 6 months when stored at 30 °C and 37 °C respectively. Stability of the heterologous protein remains unaffected during the process of heat-inactivation and lyophilization. The mass of lyophilized yeast powder remains almost unchanged during the entire period of storage. Expressed protein remains intact even after two cycles of freeze and thaws. The protease deficient strain appears ideal for the development of whole recombinant yeast-based vaccines. The cellular abundance of expressed antigen in dry powder after a year was comparable to freshly lyophilized cells. SEM microscopy showed the intact nature of cells in powdered form even after a year of storage at 30 °C. Observation made in this study showed that freeze-dry yeast powder can play a vital role in the development of thermostable vaccines.

## Introduction

In the last century, vaccines have proved to be one of the most important medical interventions in the fight against infectious diseases. Eradication of smallpox in 1980 (reviewed in **Fenner 1982)** and soon polio (reviewed in **Norrby *et al.* 2017)** are important success stories associated with the benefits of vaccines in public health. Widespread application of vaccines also leads to a sharp decline in newer cases of tuberculosis, hepatitis, measles, tetanus and other infectious diseases (**Versteeg *et al.* 2019)**. The importance of vaccines can also be underscored, by the fact that there is a race for vaccine development against novel coronavirus (SARS-CoV-2) responsible for present COVID-19 pandemic **(Callaway 2020)**. As per WHO estimates, every year, around 2-3 million lives are saved by vaccine application **(Cruz-Resendiz *et al.* 2020).** Still, millions of individuals (out of which majority are children below 5, years of age) die from vaccine-preventable diseases. Despite the significant efforts at every step, millions of people are left unvaccinated, thus putting them at the risk of getting the infection at some point in life. According to GVPA, 2011-2020 review report approximately 19.4 million infants did not receive lifesaving vaccines in 2018 (https://www.who.int/immunization/globalvaccine_action_plan/en/). Among the various issues, associated with presently licensed vaccines (summarized by **Kumar and Kumar 2019),** inherent thermolabile nature of these vaccines makes their storage, transportation and dissemination a daunting task. In the field of vaccines, this is commonly known, as the “Cold Chain” problem and, at every step, maintenance of refrigerated condition (2-8 °C in general) is a must. In many instances, maintenance of continuous cold chain raises the cost of the vaccines by about 80 % (**Bandau *et al.* 2003; Das 2004**). Exposure of vaccine to sub-optimal temperature even for short duration leads to vaccine degradation and dramatic loss in efficacy or potency which forces almost 50 % of the vaccines to be discarded before their application (**Brandau *et al.* 2003; Hill *et al.* 2016**). This scenario becomes even more relevant in resource-poor settings in countries of Asia and Africa, which lack access to vaccines **(Chen and Kristensen 2009; Das 2004).** Therefore, improving the thermostability of available vaccines at ambient temperature is highly desirable.

To address, the problem of poor thermostability and short shelf life of vaccines at ambient temperature, different approaches have been taken in the past. In the case of human enterovirus type 71, biomineralization of virus particle improves thermal stability significantly **(Wang *et al.* 2013)**. In another common approach, modification in a liquid formulation was found encouraging. For example, the addition of stabilizers like deuterium oxide, proteins, MgCl_2_, and non-reducing sugars in vaccine formulation improves their thermal stability **(Milsstein *et al.* 1997; Alcock *et al.* 2010).** Similarly, the addition of anionic nanogold particles and PEG (<u>p</u>oly<u>e</u>thene <u>g</u>lycol) improve the thermal stability of some of the vaccines significantly **(Pelliccia *et al.* 2016)**. Recently the coating of bacterial cells or viral particles in a thin film of sugar gave promising results **(Leung *et al.* 2019; Bajrovic *et al.* 2020).** All the above-highlighted approaches used the addition of one or more chemicals in vaccine preparation which, necessitating additional safety tests and other regulatory procedures. Moreover, the application of all the aforementioned approaches kept the vaccine stable only for a short duration depending upon vaccine and storage temperature. Use of these approaches makes the entire process lengthier and more cumbersome. Therefore, a procedure, which is simple, cost-effective, safe, while improving the thermal stability and shelf life of a vaccine at ambient temperature will be desirable.

It is well known and established, that the freeze-drying or lyophilization of biomolecules and biopharmaceuticals (proteins, lipids, nucleic acids) improves their stability as well as shelf life significantly **(Emami *et al.* 2018; Liao *et al.* 2004; Song *et al.* 2017).** Lyophilization has been shown to improve the stability of whole cells (example bacterial cell) and viral particles that are regularly used in conventional vaccines preparation **(Wang *et al.* 2012; Maa *et al.* 2004; Garmise *et al.* 2007)**. Although purified protein antigen can also be lyophilized and can be used as vaccine but poor immunogenicity and fast body clearance of pure protein remains an important concern. Apart from that application of purified antigen (subunit vaccines) will require use of adjuvant and will involve protein purification which will further increase cost, time and will affect the number of doses that can be available. On top of that lyophilized protein antigen will again require cold chain (**reviewed by Vartak *et al.* 2016; Kumru *et al.* 2014**). The natural adjuvant nature of yeast cell walls makes it possible to use lyophilized recombinant cells without the addition of an external adjuvant **(Stubbs *et al.* 2001)**. Unlike subunit vaccine, the yeast-based vaccines do not require protein purification. Yeast cells are efficiently taken up, by APCs **(Xiang *et al.* 2006)**. Handling of recombinant yeast is much easy and safer compared to infectious biological entities. Most importantly, the application of inactivated yeast cells is found safe and well-tolerated in human subjects **(Gaggar *et al.* 2014)**.

This forced us to investigate, whether whole recombinant yeast lyophilized powder can keep the heterologous protein (acting as an immunogen) intact when stored, at ambient temperature. In this proof of concept study, we showed that protein antigen remains stable in lyophilized yeast powder for more than a year when stored at 30 °C. Stability of heterologous fusion protein remains unaffected during the process of heat-inactivation and lyophilization. Apart from this, the mass of lyophilized yeast powder remains essentially unchanged. The observations from this study will help in developing a thermostable vaccine with long shelf life even under non-refrigerator condition (2-8 °C) using a yeast-based platform.

## Materials and methods

### Yeast strains

Haploid auxotrophic PPY12h (*arg4 his4*) **(Gould *et al.* 1992)** and protease deficient SMD1163 (*pep4 prb1 his4*) **(Gleeson *et al.* 1998)** *Pichia pastoris* strains, were used in the entire study.

### Media

YPAD media (1 % yeast extract, 2 % peptone, 0.05% adenine and 2 % dextrose), SD+CSM-His (0.17 % YNB without amino acid and ammonium sulfate, 0.5 % ammonium sulfate, 0.08 % CSM-His, 2 % agar, 2 % dextrose) **(Kumar 2019)**.

### Cloning of *E. coli* curli protein

Synthetic construct coding for *E. coli* curli was synthesized by a commercial vendor (Genewiz, New Jersey, USA) into a pUC57-Amp vector. Fragment coding of curli ORF was excised from the vector and subcloned into *Pichia pastoris* integrating vector pIB2 **(Sears *et al.* 1998)** into which cycle 3 GFP, was cloned previously **(Crameri *et al.* 1996)**. The sequence of ORF was the same as that of the original construct for which sequence, was deposited in Genebank with the following accession number MH264502 **(Kumar 2018)**. Expression of the CSGA-GFP fusion protein was under the constitutive GAP promoter (<u>G</u>lycer<u>a</u>ldehydes-3 <u>p</u>hosphate dehydrogenase). The combined mass of the fusion protein was 42.9 kDa. It is, important to note that no codon optimization for the construct was used in this study.

### Yeast transformation

*Pichia pastoris* (now officially *Komagataella phaffii* or *K. phaffii*) **(Kurtzman 2009; Kumar *et al.* 2020)** transformation was performed as described previously **(Kumar 2019)** and briefly mentioned here. The final plasmid (pRK10) was linearized by digestion with *EcoNI* which cleaves within the *HIS4* marker gene. Linearized plasmid was, transformed into PPY12h, and SMD1163 strain using the electroporation method (GenePulser Xcell from Biorad). Transformants were selected, on *His*^−^ plates and positive transformants were confirmed, both by the fluorescent microscopy and western blot.

### Protein extraction from regular cycling cells

Presence of *E. coli* CSGA-GFP protein in *P. pastoris* was confirmed by detecting CSGA-GFP fusion protein using polyclonal rabbit anti-GFP antibodies (from Life Technologies USA; cat # A-11122). Amount of protein was normalized, based on either the number of cells or by dry mass of cells. Cells were treated, with 12.5 % TCA and samples were incubated at −80 °C for one hour or overnight. On completion of incubation, samples were thawed, at room temperature and, the samples were centrifuged, at 12000 g for 8 min. The supernatant was, discarded and, the pellet was, washed twice with 80 % chilled acetone. Finally, protein pellet was air-dried and was resuspended, in 150 μL 1 % SDS and 0.2 N NaOH. Then 150 μL 2X dye (100 mM Tris HCl pH6.8, 200 mM DTT, 4 % SDS, 0.2 % bromophenol blue, 20 % glycerol) was added to the sample and samples were heated for 5 min at 95 °C using dry heating block. Samples were cooled down, to room temperature, vortexed, spanned and, an equal amount or volume of samples was loaded, in each well of 10 % SDS-PAGE along with pre-stained protein marker (Biorad, cat # 161-0376). Samples were run at constant 100V till the dye front reaches the bottom of the gel **(Kumar 2019)**.

### Western blot

On completion of SDS-PAGE run, proteins were transferred onto nitrocellulose membrane by wet transfer (at constant 100 V for 1 hour at 4 °C) as described elsewhere **(Towbin *et al.* 1979)**. The blotting membrane was incubated in blocking buffer (5 % nonfat skimmed milk powder in TBST) (TBST, 137 mM NaCl, 2.7 mM KCl, 19 mM Tris base, 0.1 % Tween 20) for 1 hour and then incubated with anti-GFP antibodies overnight at 4 °C under gentle shaking condition. Primary antibodies were removed and, the membrane was washed thrice with TBST. The membrane was again incubated with IRDye®800CW goat anti-rabbit secondary antibodies (from LI-COR, USA; cat # 926-32211) for one hour at room temperature. The membrane was again washed thrice with TBST and blot was scanned using LI-COR Odyssey software (from LI-COR, Nebraska, USA) as per manufacturer instructions.

### Heat inactivation of recombinant *P. pastoris*

A single colony of recombinant *P. pastoris,* expressing *E. coli* CSGA-GFP under GAP promoter was inoculated into 5 mL of YPAD tube. The tube was incubated, at 30 °C, 230 rpm for overnight growth (source of inoculum). The overnight grew culture was used for the inoculation of 500 mL YPAD in 2.8 L flasks. The flasks were incubated, at 30 °C, 230 rpm for 48 hours. Cells were pelleted by centrifugation at 3000 *g* for 5 minutes. Cells were washed twice with sterile water. Finally, the cell pellet was, resuspended in 25 mL sterile water and tube was incubated in a water bath maintained at 56 °C for 90 min for heat-inactivation of yeast cells **(Haller *et al.* 2007).** After 90 min, tubes were taken out of a water bath, cooled to room temperature. A small volume (10 μL) of cell suspension was taken out and plated on YPAD plate to check for the presence of viable cells. The remaining cell suspension was, centrifuged at 3000 *g f*or 5 min, the supernatant was, discarded and, pellet in tubes was put for lyophilization as described below.

### Lyophilization (freeze-drying) of whole recombinant *P. pastoris*

Recombinant *Pichia pastoris* was lyophilized as described elsewhere **(Patterson *et al.* 2015)** and briefly mentioned here. Cells were harvested in 50 mL falcon tube by centrifugation at 3000 g for 5 minutes and removal of supernatant. Lyophilization was performed on the cell pellet using the AdVantage 2.0 Bench Top Freeze Dryer/Lyophilizer (from SP Scientific). Samples in tubes were regularly checked, for the formation of lyophilized powder. On completion of lyophilization, tubes were, taken off and the combined mass of tube and lyophilized yeast powder was noted.

### Storage of freeze-dried yeast powder

On completion of the process of freeze-drying, tubes were removed from the lyophilizer and caps were tightly closed. Tubes were kept in cardboard boxes and stored in a separate incubator (in the dark away from direct sunlight, no humidity) operating at 30 °C and 37 °C. Then the weight of tubes was noted, at a regular interval. All steps were performed under normal lab environment (i.e. not in an inert and sterile atmosphere).

### Protein extraction from lyophilized powder

A known amount of lyophilized yeast powder was, taken in a sterile Eppendorf tube. The powder was resuspended in 200 uL of 12.5 % TCA and stored at −80 °C for 1 hour or overnight. The subsequent procedure is the same as described above for protein extraction.

### Freeze and thawing of lyophilized yeast powder

As above, a known amount of lyophilized yeast powder was, taken into fresh sterile Eppendorf tubes. Tubes were, stored at −20 °C and −80 °C for, two hours followed by incubating the tubes at 30 °C for two hours. This process was repeated, depending on the number of freeze-thaw cycles. A known amount of yeast powder was also taken and incubated at 30 °C as a control. Protein extraction was performed as described above.

### Field Emission Scanning Electron Microscopy

The shape and surface morphology of lyophilized yeast cells stored at 30 °C for a year was analyzed, by Field Emission Scanning Electron Microscopy (ZEISS Gemini Sigma 500 VP, Carl Zeiss Microscopy LLC, NY, USA). Samples were prepared, by mounting a small amount of lyophilized yeast powder on a double-sided carbon tape on a metal SEM stub. Gently spread the yeast powder across the surface of the tape. Compressed air was, used to remove loose lyophilized yeast cells. Yeast samples were sputter-coated with a 3 nm thin layer of Gold-Palladium for ensuring the conductivity for electrons beam. A beam strength of 5.0 kV and a working distance in the range of 8–9 mm was, used to visualize yeast samples.

### Fluorescence Microscopy

Expression of *E. coli CSGA-GFP* fusion protein in both SMD116 and PPY12h *Pichia pastoris* was confirmed, by fluorescence microscopy. For each strain, 100 uL cycling cells, were taken into sterile Eppendorf tube, cells were pellet down by centrifugation at 3000 g for 3 min. The supernatant was discarded, and the cell pellet was resuspended, in 1 mL sterile Milli Q water. 5 uL of cell suspension was transferred, onto a glass slide. Cells were fixed by adding an equivalent volume of 1 % agarose. Images were captured, using plan apochromat 100× 1.40-NA oil immersion objective on a motorized fluorescence microscope (Axioskop 2 MOT plus; Carl Zeiss) coupled to a monochrome digital camera (AxioCam MRm; Carl Zeiss). Images analysis was performed, using Axio vision software **(Kumar 2019)**.

## Results

### The basic assumption behind the study

In our previous study, we showed that stationary phase yeast cells were able to keep the heterologous protein intact when stored at room temperature (23-25°C) for a year **(Kumar,2018)**. Unfortunately, the gradual degradation of cells over time suggests that long-term storage of protein antigen even within cells in a liquid environment is not feasible **(Kumar 2018)** Therefore, we looked at other ways for long-term storage of cells at ambient temperature. It is a common observation that active dry yeast powder sold commercially is stable (a significant proportion of cells retain viability) for almost two years when stored under non-refrigerated conditions (at 23-25 °C). Even under recommended conditions of storage and handling most of the available vaccines remain potent for up to 1.5 to 3 years. Therefore it raised the possibility that dry yeast powder can act as a simple yet an important, means for long-term storage of immunogen (vaccine) under room temperature (ideally around two years) **(Clenet 2018; Chen *et al.* 2010; Galazka *et al.* 1998; Chen and Zehrung 2013)**. Through this proof of concept study, we tested whether, lyophilized yeast cells can be used, to store protein immunogen at ambient for year or more. For this, we expressed *E. coli* curli-GFP fusion protein (*CSGA-GFP*) in *P. pastoris*. Curli is a bacterial surface protein (fimbri) involved in cell adhesion and biofilm formation **(Barnhart *et al.* 2006; Nguyen *et al.* 2014)**. Curli is capable of raising immune response, thereby acting as a good immunogen (**Barnhart *et al.* 2006**) and a suitable, candidate for the proof of concept study.

### Expression of *E. coli CSGA-GFP* in *Pichia pastoris*

*E. coli* curli was subcloned into *P. pastoris* integrating vector pIB2 as described in the materials and methods section. Cartoon presentation of the final plasmid used for the expression of *E. coli CSGA-GFP* fusion protein is shown, in Figure 1A. For checking the expression of the CSGA-GFP fusion protein, we randomly selected eight colonies from the transformant plate and patched them on a fresh selection plate (CSM-HIS plate). Next day, the expression of the fusion protein was first confirmed, by fluorescent microscopy (Figure 1B for PPY12h background). Later in a separate experiment, the same plasmid was transformed, into SMD1163 strain. Integration of plasmid and expression of CSGA-GFP was confirmed by fluorescent microscopy (Figure 1C).

**Figure 1.**
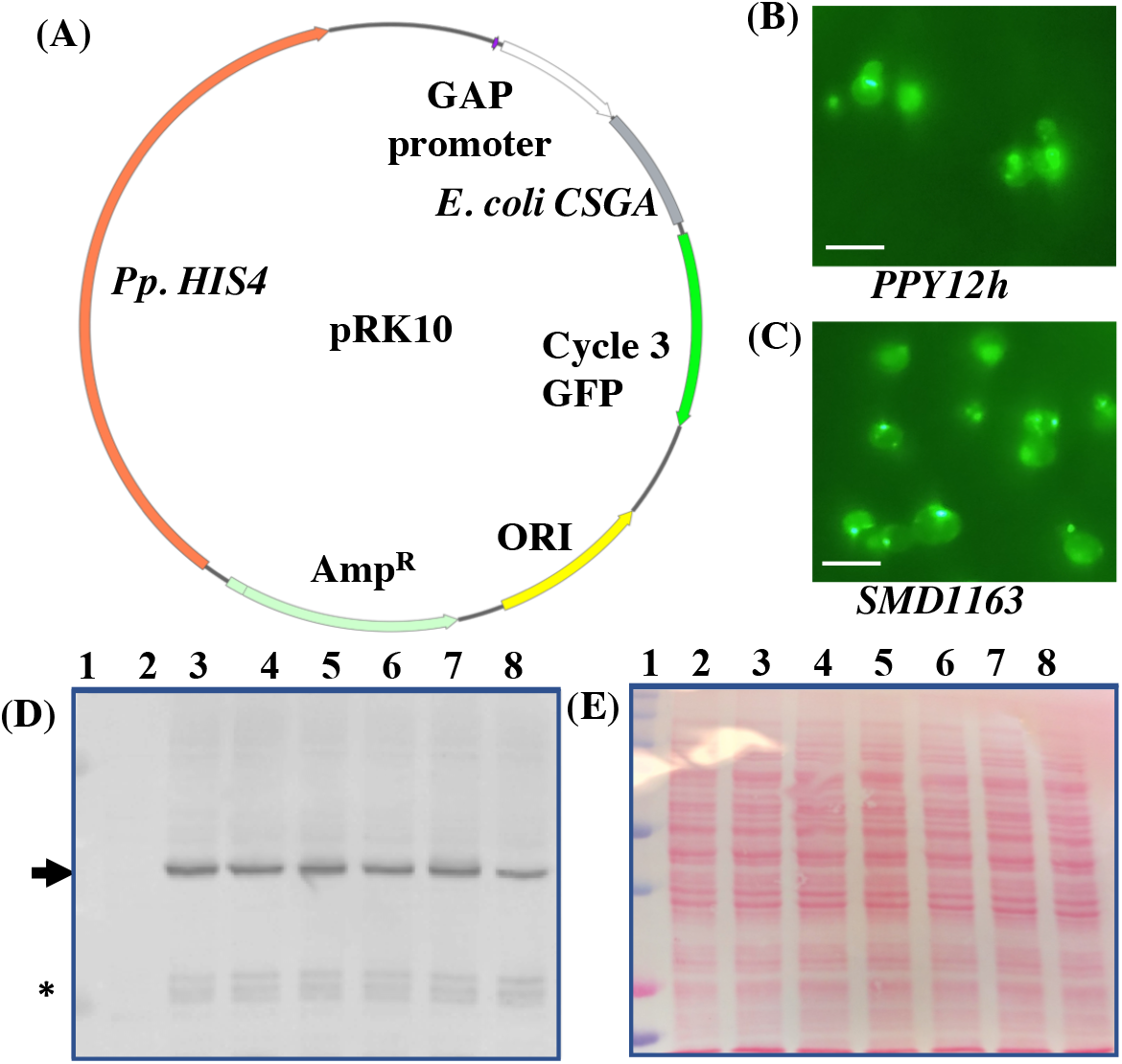
Expression of *E. coli* CSGA-GFP fusion protein in *P. pastoris*. (A) Map of the final plasmid used for expression of the CSGA-GFP fusion protein. Expression of a CSGA-GFP fusion protein in (B) PPY12h and (C) SMD1163 strain. Scale bar represents 5 uM. (D) Image of western blot showing expression of the CSGA-GFP fusion protein in PPY12h. Bands of interest are pointed by arrows towards them, while free GFP is pointed by an Asterisk mark. (E) Proper loading and protein transfer are shown, by Ponceau S stained blot image. Well, 1 (pre-stained protein marker), well 2 (empty vector as a negative), well 3-8 (transformants).

Expression of the fusion protein, CSGA-GFP, was further confirmed by western blot. 2 OD_600nm_ of cells were used for protein extraction for western blot. Blot image in Figure 1D shows that all the selected colonies were positive. Ponceau-S stain image of the same blot is shown, as a loading control (Figure 1E) which confirm the proper loading of protein in each well. Specific anticipated bands are highlighted, by an arrow pointing towards them. We do not get any band in empty vector control strain used as a negative control. The absence of any signal from negative control shows the specificity of GFP antibodies used in the experiment. The expression of the heterologous protein was endogenous. Overall we can say that we were able to express the fusion protein in *P. pastoris.*

### Heat inactivation and lyophilization do not affect the stability of expressed antigen

Yeast species especially *S. cerevisiae* and *P. pastoris* are non-pathogenic and are in the list of GARS and routinely used for the production of diverse biomolecules for human consumption **(Ramchuran *et al.* 2005; Basanta *et al.* 2010)**. Even then. direct administration of live recombinant yeast cells into human subjects is not advisable from the point of safety. However, the level of immune response mounted on the application of whole recombinant yeast is independent of live or dead nature of yeast **(Lu *et al.* 2004; Franzusoff *et al.* 2005)** raising the possibility for application of inactivated recombinant yeast. Heat-inactivation of yeast is rapid, simple and more convenient compared to chemical based-inactivation of bacterial or viral particles for vaccine preparation. The combined effect of heat-inactivation and lyophilization on the stability of heterologous protein was missing and through this study, we tried to fill that gap. Heat inactivation of yeast cells was performed as described in the materials and methods. Heat inactivation was confirmed by plating small volume of cell suspension on YPAD plates (Figure 2B) along with untreated control cycling cells (Figure 1A). Our present western blot data showed that the stability of heterologous proteins remains essentially unaffected during heat-inactivation (Figure 2C) and heat-inactivation followed by lyophilization (Figure 2E). Proper loading and transfer of protein are shown by Ponceau-S stained blots (Figure 2D and 2E respectively). Protein amount was normalized, by the weight of cells used for protein extraction. Results of Figure 2C are in accordance with our previous study **(Kumar 2018)**. Based on the present data, it can be said that together heat-inactivation and lyophilization have no affect protein stability.

**Figure 2.**
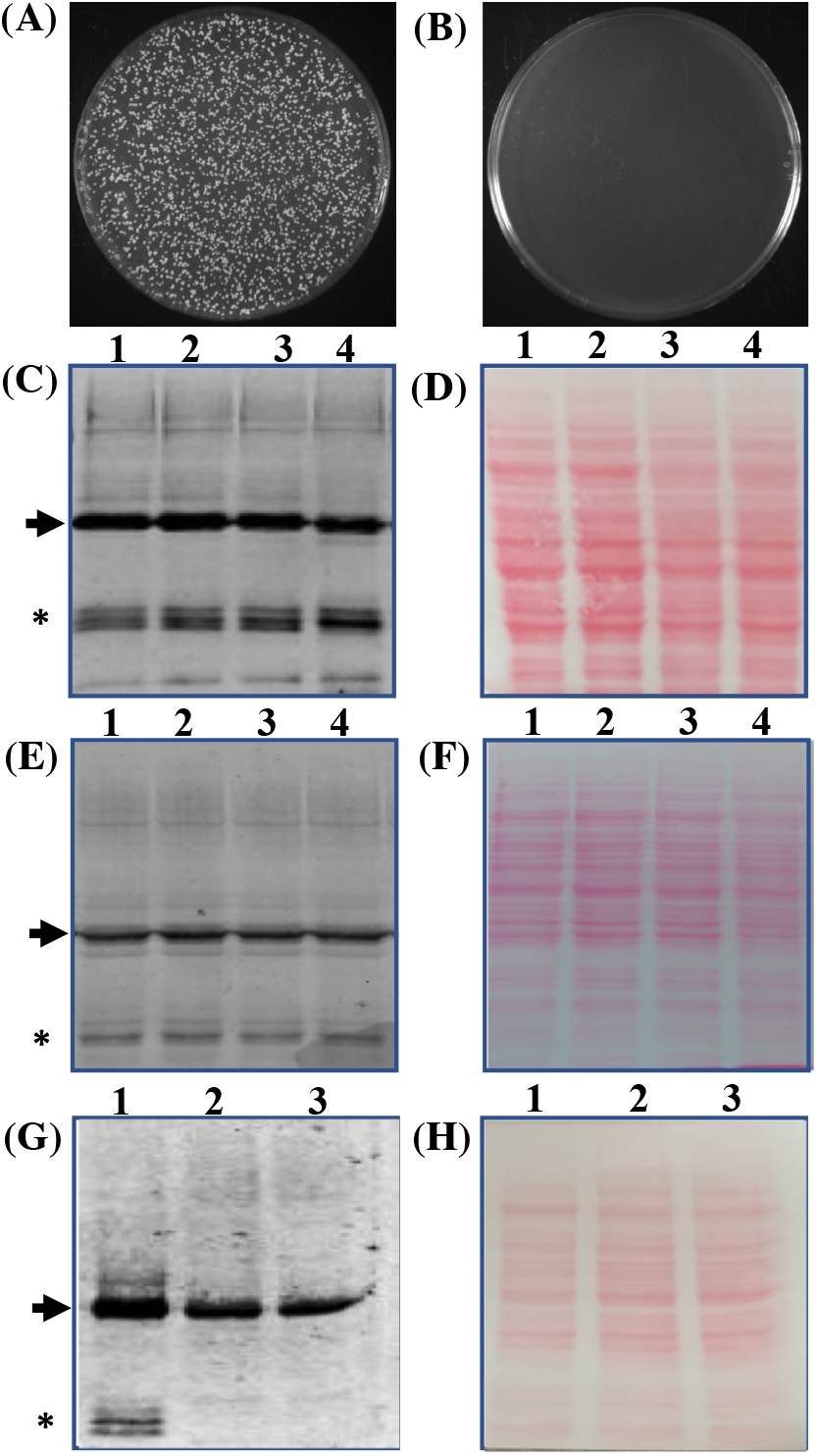
Heat-inactivation and freeze-drying do not affect protein stability. (A) Control untreated viable cells. (B) Complete loss of viability on heat-inactivation. (C) Western blot image showing the effect of heat-inactivation on protein stability. Well, 1, 2 (untreated cycling cells); well 3 4 (heat-inactivated cells). (D) Proper loading and protein transfer is shown by Ponceau S stain blot image. Western blot image showing the effect of freeze-drying on protein stability (E) well 1, 2 (control cycling cells); well 3,4 (lyophilized yeast cells). (F) Proper loading and protein transfer is shown by Ponceau S stain blot image. Bottom free GFP bands are due to vacuolar protease (G) well 1 (PPY12 strain); well 2-3 (SMD1163, protease deficient strain, lack Prb1 cytosolic and Pep4 vacuolar protease). (H) Proper loading and protein transfer is shown by Ponceau S stain blot image. Bands of interest are pointed by arrows towards them, while free GFP is pointed by an Asterisk mark.

### The protease deficient yeast strain improves the stability of the expressed fusion protein

In our previous section, we showed that process of heat-inactivation and heat-inactivation followed by lyophilization does not affect the stability of CSGA-GFP in yeast cells. But we do see a very prominent band around 25 kDa region of the blot. Whether the observed bands were due to degradation of the fusion protein on heat-inactivation and lyophilization, or whether they were a result of protease activities in cytosol and vacuoles was not clear. To sort out this, we transform the same construct into protease deficient strain (SMD1163), which lacks Prb1 (cytosolic) and Pep4 (vacuolar). Our present western blot data pointed towards the idea that observed lower bands are the result of vacuolar protease action (Figure 2G). Like previous blots, we again observed lower bands in PPY12h strain used as a control (Figure 2G, well1), but failed to detect the same bands in SMD1163 (Figure 2G, well 2,3). The presence or absence of lower bands in PPY12h and SMD1163 respectively is not due to the difference in the amount of protein loaded into each well was confirmed by Ponceau S stained blot image (Figure 2H). Therefore, it can be concluded that lower observed bands are not due to heat-inactivation or lyophilization but may be due to the action of cellular proteases. Our observation is supported by the fact that, during stress, autophagic pathways get activated, which forces the degradation of cellular components in vacuoles or lysosomes **(Klionsky *et al.* 2007; Takeshige *et al.* 1992; Kumar *et al.* 2020)**. Therefore lower bands detected in PPY12h and their absence in SMD1163 may be due to autophagic degradation of fusion proteins in vacuoles and GFP (26.7 kD), which is quite stable in vacuoles (whose free release in the vacuole is used in autophagic assays) was detected by antibodies **(Klionsky *et al.* 2007; Takeshige *et al.* 1992)**.

### Mass of freeze-dried yeast powder remains unchanged during storage

After confirming that heat-inactivation, as well as freeze-drying, do not affect the stability of expressed protein, we increase the volume of culture to 1 L. PPY12h strain expressing CSGA-GFP was grown in bulk volume (1 L). Cells were, harvested heat-inactivated and freeze-dried as described in Materials and Methods. After lyophilization, tubes were taken off from lyophilizer, screwed the cap and weighed. Initial mass was taken and stored at 30 °C for one year. The weight of the tube was, then checked, regularly and sometimes the tubes were opened, for a short time. Table 1 is showing the data for the mass of powder stored at 30 °C. Similarly, we checked, the weight of freeze-dried yeast powder stored at 37 °C for six months (Table 2). Present data showed that mass of freeze-dried powder remains essentially unchanged under both conditions of storage. The slight variation observed in weight may be due to moisture that might be entered into tubes when tubes were open for some time. Note in Table 1 data is shown for two conditions. In one condition cells were lyophilized after-heat inactivation (Table 1 second and third column) and in another, cells were lyophilized without heat-inactivation (table 1 fourth and fifth column). Based on the data shown in tables, it can be said, that the mass of stored lyophilized cells does not change significantly.

**Table 1.** Mass of lyophilized yeast powder stored at 30 °C for one year.

| *Month* | *Heat inactivated* |  | *Without heat inactivated* |  |
| --- | --- | --- | --- | --- |
|  | *Tube 1* | *Tube 2* | *Tube 1* | *Tube 2* |
| **0** | 16.88 | 16.97 | 17.06 | 17.26 |
| **2** | 16.97 | 16.96 | 17.14 | 17.29 |
| **4** | 16.95 | 16.91 | 17.14 | 17.27 |
| **6** | 16.98 | 16.93 | 17.13 | 17.26 |
| **8** | 16.95 | 16.91 | 17.15 | 17.21 |
| **10** | 16.96 | 16.93 | 17.16 | 17.21 |
| **12** | 16.97 | 16.95 | 17.15 | 17.22 |

**Note:** Net mass is in grams. Reported mass is the combined mass of tube and powder.

**Table 2.**
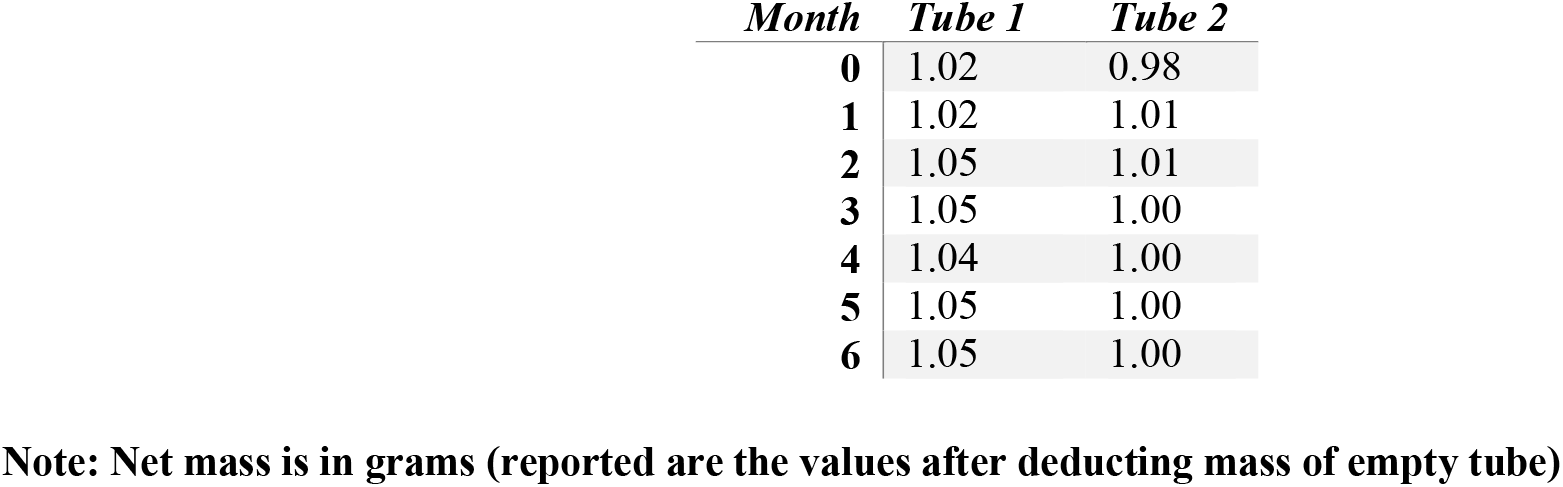
Mass of lyophilized yeast powder stored at 37 °C for six months.

### Cells in lyophilized powder retain their intactness

Next we asked whether cells in lyophilized powder also remain intact by checking the morphology and surface appearance of cells using SEM microscopy. A small amount of lyophilized powder (figure 3A cells were lyophilized without heat-inactivation and, figure 3B cells were lyophilized after heat-inactivation) was taken and SEM microscopy was performed. SEM images showed that the cells remain intact in lyophilized powdered form when stored at 30 °C for more than a year (figure 3C and 3D). Although cells were intact under both the condition, we observed a difference in the texture of the powder. Freeze dry powder of cells without heat-inactivation appears less compact and less dense (figure 3A) whereas heat-inactivated powder appear more compact and denser (figure 3B). Thus, it can be said, that cells in lyophilized powdered form remain intact for a year even when stored under ambient or room temperature.

**Figure 3.**
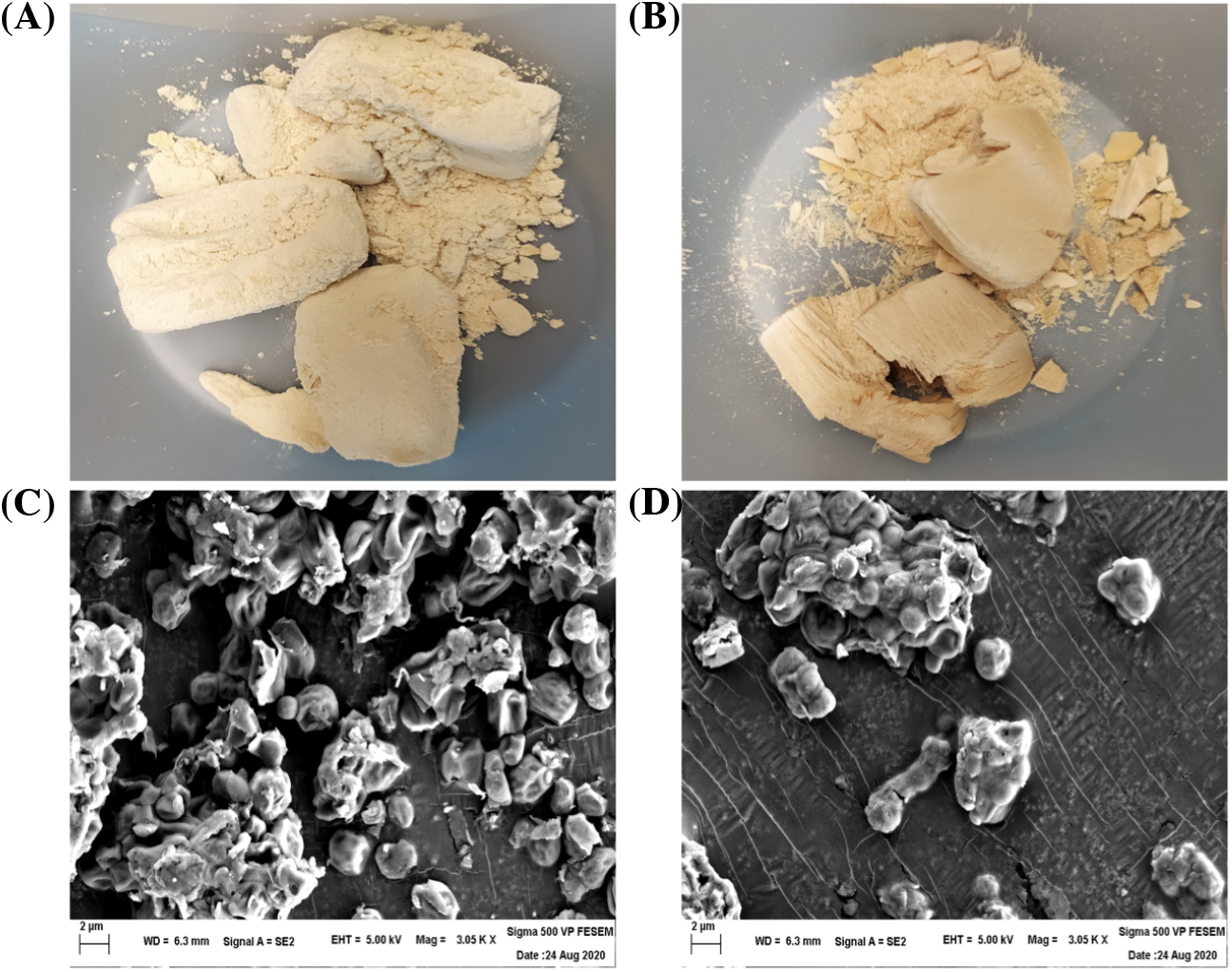
Cells in lyophilized powder remain intact for more than a year. Image of lyophilized yeast powder stored at 30 °C for more than a year (A) cells was lyophilized without heat-inactivation and (B) cells were heat-inactivated before lyophilization. SEM images showing external morphology of lyophilized cells after one year (C) lyophilized cells without heat-inactivation and (D) lyophilized cells after heat-inactivated.

### Expressed antigen remains stable in lyophilized yeast powder stored at 30 °C temperature

Although SEM data in the previous section confirmed intact nature of lyophilized yeast cells after a year of storage at 30 °C, whether expressed protein also remained intact was unknown. To check the stability of the expressed protein, we took equal amounts of lyophilized powder from both the condition and extracted the protein. An equal amount of whole cell lysate was loaded on 10 % SDS-PAGE and, the fusion protein was detected using anti-GFP antibodies. Our present western blot data showed that expressed protein remains stable in powdered yeast stored at 30 °C for a year (figure 4). The expressed protein remains stable under both the conditions in which cells were lyophilized without heat-inactivation (figure 4A) and in which cells were lyophilized after heat-inactivation (figure 4B). Before incubation in blocking buffer, blots were stained with Ponceau S for showing the loading control (figure 5C and 5D for figure 4A and 4B respectively). Encouraged from the stability of the protein in lyophilized powder stored at 30 °C for a year, we checked whether expressed protein also remains stable in lyophilized yeast cells stored, at 37 °C for six months. Our present western blot data showed that expressed protein remains stable in powdered yeast even when stored at 37 °C at least for six months (figure 4E). Loading control is shown in figure 4F. Therefore, it can be, concluded that expressed protein remains stable in lyophilized yeast cells when stored at a temperature ranging from 30 °C −37 °C for quite a long time.

**Figure 4.**
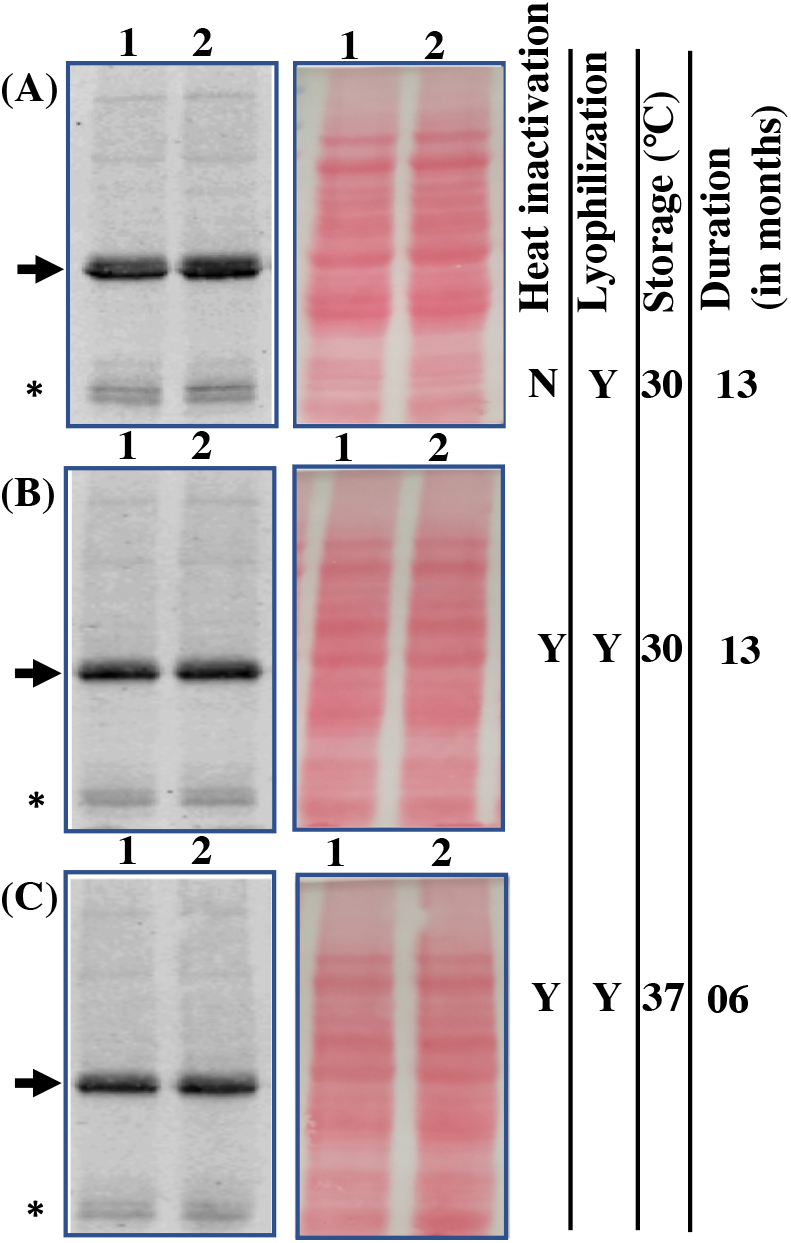
Expressed protein remains stable in powdered yeast. (A) Stability of CSGA-GFP in freeze-dried yeast powder stored at 30 °C after one year. (B) Stability of CSGA-GFP in heat-inactivated lyophilized yeast powder stored at 30 °C after one year. (C) Stability of heat-inactivated lyophilized yeast powder stored at 37 °C after six months. In each case, proper loading is shown, by Ponceau S stained blot image. Note, the protein amount was normalized, based on yeast powder used for protein extraction. Bands of interest are pointed by arrows towards them, while free GFP is pointed by an Asterisk mark. Note in figure Y stand for yes and N for no for condition mentioned in the figure.

**Figure 5.**
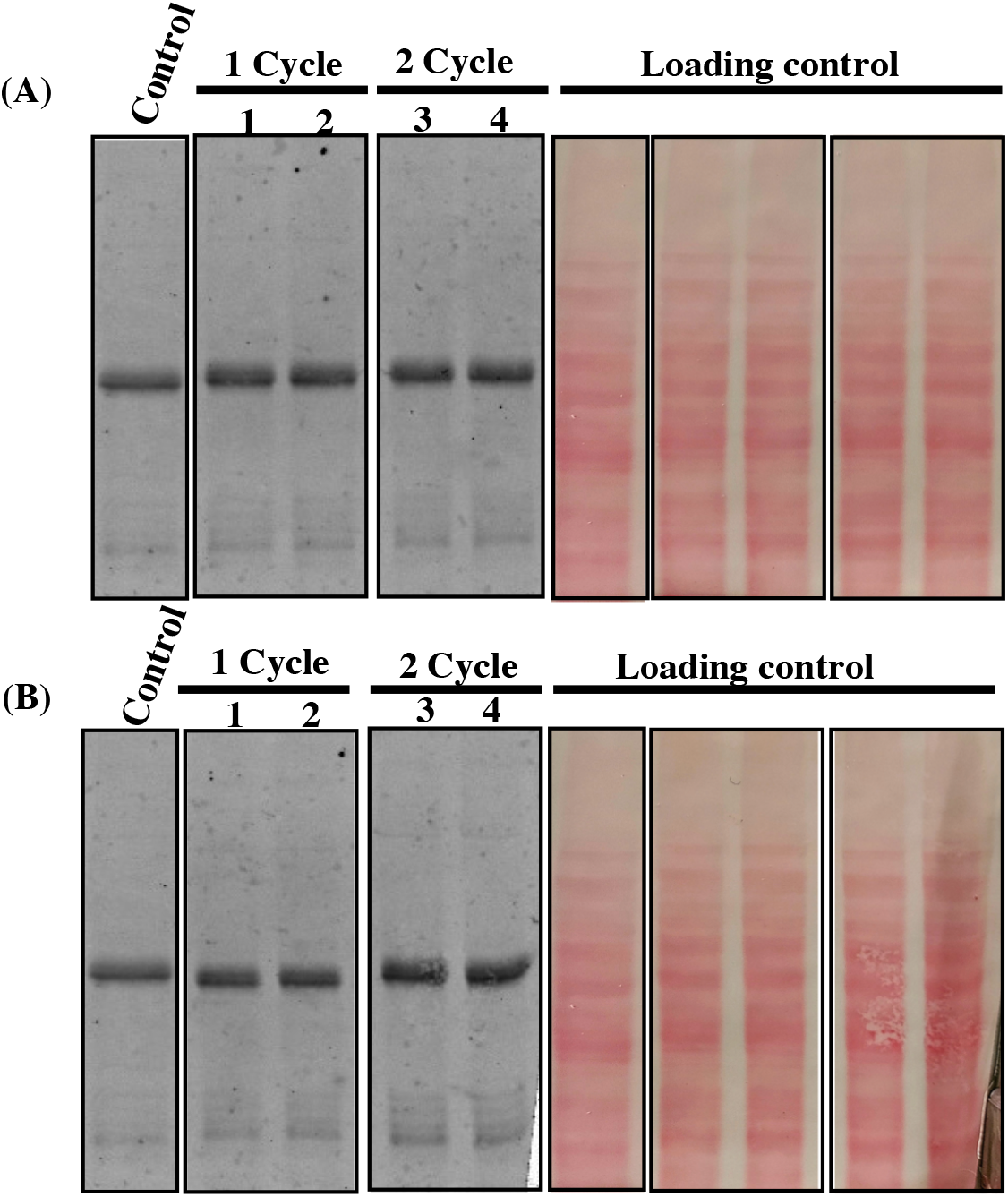
Freeze and thaws do not affect the stability of the expressed protein. (A) Effect of freeze and thaw at −20 °C. Control (lyophilized powder stored at 30 °C), well 1,2 (once the cycle of freeze and thaw), well 3,4 (two-cycle of freeze and thaw). (B) Effect of freeze and thaw at −80 °C. Same common control as above, well 1,2 (one cycle of freeze and thaw), well 3,4 (two-cycle of freeze and thaw). Note, in all case the protein amount was normalized based on the weight of powder and, each sample was taken in duplex. Loading and proper transfer are shown through Ponceau S stained of same blots. Also, a control sample is common in panel A and B.

### Freeze and thaw do not affect protein stability in powdered yeast

Most of the commonly used vaccines are stored at 2-8 °C, while some of them are also stored at −15 °C to −50 °C (e.g. <u>m</u>easles, <u>m</u>umps and <u>r</u>ubella or MMR_I_ is stored at +8 °C to −50 °C, https://www.merckvaccines.com/mmr/storage-handling/). However for most vaccines, exposure to sub-optimal temperature (i.e. more than 2-8 C or below freezing temperature) even for a short duration is known to reduce vaccine potency dramatically) **(Brandau *et al.* 2003; Hill *et al.* 2016)**. An ideal vaccine should remain stable and retain potency even when stored under the non-refrigerated condition as well as under accidental freeze condition. For investigating the effect of freeze and thaw on the stability of the heterologous protein in lyophilized yeast cells, known amount of yeast powder was taken into separate tubes. Tubes were stored at −20 °C and −80 °C for two hours. Freeze and thaw were performed in one cycle and two-cycle. For each condition, samples were taken, in duplex. After a freeze, thawing was performed by incubating samples at 30 °C for two hours. Our present western blot data showed that expressed protein in lyophilized yeast powder remain stable following one as well as two cycles of freeze and thaw at both −20 °C (figure 5A) and −80 °C (figure 5B). Ponceau S stained image of the membrane (as a loading control), is shown next to the blot. A known amount of yeast powder, stored at 30 °C separately was taken, as a control. Conditions of freeze and thaw are mentioned, in the materials and methods section. Based on present data, it can be said that expressed protein remains stable both under the non-refrigerated condition for a year as well as under freeze and thaw condition.

### The abundance of expressed protein after a year was similar to fresh cells

In the previous section, we showed that the expressed CSGA-GFP fusion protein was stable in lyophilized yeast powder when stored at 30 °C for a year and also survived two cycles of freeze and thaw. But whether the expressed protein deteriorated over time and to what extent still needed to be determined. To answer this question, we compared the level of expressed protein in yeast powder stored for a year to the freshly lyophilized yeast cells. Before comparing, the level of fusion protein we checked the overall protein content in the two samples (Figure 6A). Although an equal amount of whole-cell lysate, was loaded for each sample, we still see a slightly low level of proteins in a sample from year-old powder (well 6,7) compared to freshly lyophilized cells (well 4,5). Apart from this we also observed relatively more background in a year old sample. Whether the observed difference in the level of proteins and background is due to the difference in protein extraction efficiencies or protein degradation is unclear. But we do find a problem in re-suspending the powder entirely even on vigorous vortex. We also loaded an equal amount of cell lysate from normal cycling cells as control (well 2,3).

**Figure 6.**
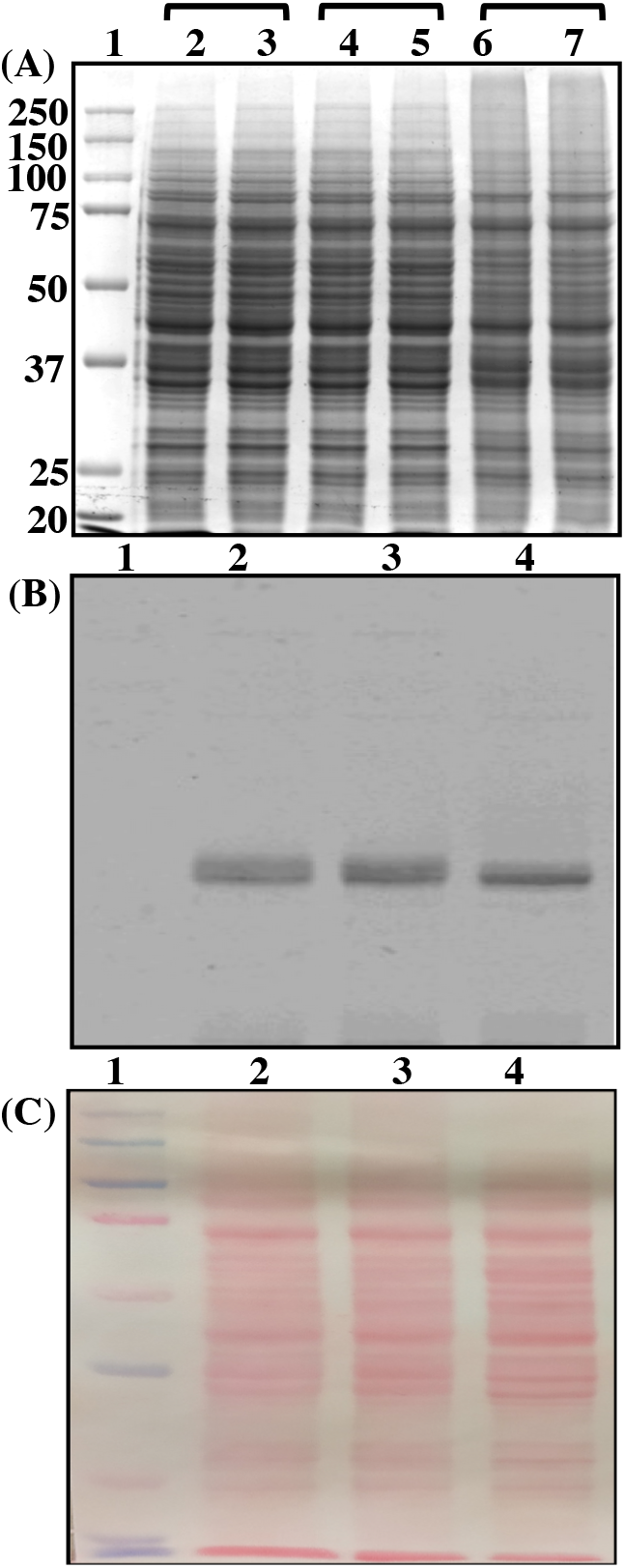
The abundance of heterologous protein in a year old powdered and freshly lyophilized cells are similar. (A) The efficiency of protein extraction from different samples. Cycling cells without lyophilized (well 2,3), an equivalent number of cells after lyophilized powder (well 4,5), an equivalent amount of lyophilized powder stored at 30 °C for one year (well 6,7), well 1 for protein marker. (B) The abundance of expressed protein is similar in freshly lyophilized cells and year-old powder lyophilized powder after one year at 30 C (well 2,3), freshly prepared powder (well 4) and protein marker (well 1). The same blot was Ponceau S stained before blocking in skimmed milk powder in TBST shown as a loading control (C).

After looking at the overall proteome of lyophilized cells stored for a year and freshly lyophilized cells, we proceed to compare the level of the expressed fusion protein in two samples. Our present data (Figure 6B) shows that level of the CSGA-GFP fusion protein is almost similar in freshly lyophilized cells (figure 6B, well 4) to that of one-year-old powder (figure 6B, well 2,3). Note we used the same sample for western blot which, were used in running SDS-PAGE shown in figure 6A. And the slight difference which appears in two samples may be due to the issues mentioned above. Loading control is shown, by Ponceau-S stained image of the same blot (figure 6C). To capture any possible difference in samples blot images were captured at low intensity of exposure (data for high exposure, is not shown).

## Discussions

Storage, transportation and distribution of vaccines before their final application are still presents difficulties. On one hand, exposure of vaccine to temperatures more than the recommended temperatures (generally 2-8 °C) leads to vaccine degradation, denaturation and finally loss of vaccine potency or efficacy **(Brandau *et al.* 2003; Hill *et al.* 2016)**. On the other hand, exposure of vaccines to below freezing conditions also affect vaccine potency **(Lloyd *et al.* 2015; Kumru *et al.* 2014)**. Therefore, maintenance of optimum conditions from point of manufacturing till the final application is required which is more difficult in less well-developed countries. A solution to this is the development of thermostable vaccines. Availability of thermostable vaccines will be an important, step in global immunization step **(Lee *et al.* 2017**). The thermostable nature of future vaccines will not only make vaccine transport, storage and distribution more convenient and economical but, will also help in saving a huge, amount of life-saving vaccines. Currently, the development of thermostable vaccines are yet unmet, with little success in which thermal stability was observed (only for a few months at best and that too in few cases) (**Leung *et al.* 2019; Chu *et al.* 2016; Mistilis *et al.* 2017; Hassett *et al.* 2013, 2015; Chen *et al.* 2010; Ohtake *et al.* 2010, 2011; Lovalenti *et al.* 2016**). By using thin-film coating, one study was able to keep the adenovirus stable at room ambient temperature for three years (**Bajrovic *et al.* 2020).** It will be interesting to see whether this approach can be applied to other vaccines. But this study showed the possibility that a vaccine may be stored and transported at ambient temperature. So far, most of the approaches used for enhancing the thermal stability and shelf life of vaccines at ambient temperature require the addition of different chemicals. Assessing the safety of each chemical added in formulation requires more regulatory clearance, time and money. Moreover, whether the developed formulation applies to a wide range of vaccines also remain a matter of future research. Unlike conventional vaccine preparation which requires growth of bacterial culture or viral particles followed by formalin-based inactivation which itself is quite lengthy, growth and heat-inactivation of yeast are rapid and more economical.

The degradation of expressed immunogen by vacuolar proteases may pose a serious issue in the development of a whole recombinant yeast-based vaccine. The degradation of immunogen will not only reduce the level of the immunogen in cells but may also affect the level of the immune response. It is, therefore, important to use protease deficient strain (like SMD1163) **(Gleeson *et al.* 1998)**. Compared to budding yeast, the post-translation modification *P. pastoris* is closer to those of higher eukaryotes **(Cregg *et al.* 1993)**. The fact that P. pastoris is better fermenter is other added advantages **(Cereghino and Cregg, 2000)** when a large amount of vaccine is needed such as during a pandemic.

Surprisingly, we do not observe any bacterial contamination in yeast powdered stored for the year even though we opened the tubes several times in a year. This can be the result of heat-inactivation followed by lyophilization. This observation showed, that lyophilized yeast powder can be, stored for a long time even in the absence of antibiotics. Several conventional vaccine preparations involve the use of antibiotics such as MMR which not only increase the cost of vaccines production but also cause an allergic response in some people **(Chung, 2014)**.

Although we have shown the data for the stability of heterologous protein for a year (at 30 °C) and six months (at 37 °C), we expect that expressed protein may remain stable for much longer (as a result we are still keeping our samples at the aforementioned condition). Further, it will be interesting to see whether recombinant yeast lyophilized powder stored at ambient for a year or more can mount a specific immune response *in vivo* models like a mouse. Previous observation that APCs can process freeze-dried yeast cells and can mount an immune response relates to our present and future study in this direction **(Patterson *et al.* 2015).** It is almost impossible to mimic the real scenario, but our present data showed that lyophilized yeast powder, can prevent immunogen degradation due accidental freeze and thaw that was lacking in other approaches used for the development of thermostable vaccines. Yeast-based vaccines, which are in different phases of clinical trials may present opportunities where the present method for developing thermostable vaccines can be adopted directly **(Kumar and Kumar, 2019; Ardiani *et al.* 2010).** Moreover, there is no obvious barrier why our approach cannot be directly adopted for subunit vaccines where protein immunogen have been already characterized **(Agwale *et al.* 2002; Zhang *et al.* 2015; Schiller and Lowy, 2015)**.

Overall, we have shown that a dry yeast powder is a simple yet effective way for long-term storage of vaccines (immunogen) under non-refrigerated condition. Data shown in this report showed that yeast-based approach for thermostable vaccine development prevents vaccine deterioration both at high temperature (above 2-8 °C) as well as below freezing condition (below 0 °C) which is common in countries of Europe, North America and parts of Asia. We believe that this cost effective freeze-dried yeast powder approach may be one solution to the cold chain problem and will go towards attaining the universal immunization program.

## List of abbreviations

SEM: Scanning electron microscope
GVAP: Global Vaccine Action Plan
SARS-CoV-2: severe acute respiratory syndrome coronavirus 2
COVID-19: coronavirus disease of 2019
APCs: antigen-presenting cells

## Acknowledgements

We are thankful to the University of California San Francisco (UCSF) San Francisco, California-USA for providing space and other necessary facilities towards completing this manuscript. We are also thankful to Prof. Geoff Lin-Cereghino from Pacific University, Stockton, California-USA for providing SMD1163 strain, Prof. Robert Stroud from UCSF for electroporation of yeast cells. We are also thankful to the staff at the University of California Berkeley Electron Microscope Laboratory for advice and assistance in electron microscopy, sample preparation and data collection. We also extends our sincere thanks to Prof. Tejal A Desai for allowing access to lyophilizer. We are specially thankful to Prof. Jennifer C. Fung for allowing us to carry out this work in her lab.

## Authors Contribution

RK conceived, designed, performed the experiments, analyzed the data and wrote the manuscript. BNK perform SEM and fluorescence microscopy.

## Declaration

Authors declare that the present study does not involve any human subjects or animal handling by any author mentioned in this study.

**For Graphical abstract**

**Schematic showing the basic workflow of the entire study.**

## References

1. Agwale SM, Shata MT, Reitz MS Jr et al. A Tat subunit vaccine confers protective immunity against the immune-modulating activity of the human immunodeficiency virus type-1 Tat protein in mice. PNAS USA 2002;99:10037–41.

2. Alcock R, Cottingham MG, Rollier CS et al. Long-Term Thermostabilization of Live Poxviral and Adenoviral Vaccine Vectors at Supraphysiological Temperatures in Carbohydrate Glass. Science Translational Medicine 2010;2:19ra12

3. Ardiani A, Higgins JP, Hodge JW. Vaccines based on whole recombinant Saccharomyces cerevisiae cells. FEMS Yeast Res. 2010;10:1060–9.

4. Bajrovic I, Schafer SC, Romanovicz DK et al. Novel technology for storage and distribution of live vaccines and other biological medicines at ambient temperature. Sci Adv 2020;6:eaau4819.

5. Barnhart MM, Chapman MR. Curli biogenesis and function. Annu Rev Microbiol. 2006;60:131–147.

6. Basanta A, Gómez-Sala B, Sánchez J et al. Use of the yeast Pichia pastoris as an expression host for secretion of enterocin L50, a leaderless two-peptide (L50A and L50B) bacteriocin from Enterococcus faecium L50. Appl Environ Microbiol. 2010;76:3314–24.

7. Brandau D, Jones L, Wiethoff C et al.. Thermal stability of vaccines. Journal of Pharmaceutical Sciences 2003;92:218–231.

8. Callaway E. The race for coronavirus vaccines: a graphical guide. Nature. 2020;580:576–577.

9. Cereghino JL, Cregg JM. Heterologous protein expression in the methylotrophic yeast Pichia pastoris. FEMS Microbiol Rev. 2000;24:45–66.

10. Chen D, Kapre S, Goel A et al. Thermostable formulations of a hepatitis B vaccine and a meningitis A polysaccharide conjugate vaccine produced by a spray drying method. Vaccine 2010;28:5093–9.

11. Chen D, Kristensen D. Opportunities and challenges of developing thermostable vaccines. Expert Rev Vaccines 2009;8:547–557.

12. Chen D, Zehrung D. Desirable attributes of vaccines for deployment in low-resource settings. J Pharm Sci 2013;102:29–33.

13. Chu LY, et al. Enhanced Stability of Inactivated Influenza Vaccine Encapsulated in Dissolving Microneedle Patches. Pharm. Res. 2016;33:868–878.

14. Chung EH. Vaccine allergies. Clin Exp Vaccine Res 2014;3:50–57.

15. Clénet D. Accurate prediction of vaccine stability under real storage conditions and during temperature excursions. Eur J Pharm Biopharm 2018;125:76–84.

16. Crameri A, Whitehorn EA, Tate E et al. Improved green fluorescent protein by molecular evolution using DNA shuffling. Nat Biotechnol 1996;14:315–9.

17. Cregg JM, Vedvick TS, Raschke WC. Recent advances in the expression of foreign genes in Pichia pastoris. Biotechnology (N Y) 1993;11:905–10.

18. Cruz-Reséndiz A, Zepeda-Cervantes J, Sampieri A et al. A self-aggregating peptide: implications for the development of thermostable vaccine candidates. BMC Biotechnol. 2020;20:1.

19. Das P. Revolutionary vaccine technology breaks the cold chain. Lancet Infect Dis 2004;4:719.

20. Emami F, Vatanara A, Park EJ et al. Drying Technologies for the Stability and Bioavailability of Biopharmaceuticals. Pharmaceutics 2018;10:131.

21. Fenner F. A successful eradication campaign. Global eradication of smallpox. Rev Infect Dis 1982;4:916–930.

22. Franzusoff A, Duke RC, King TH et al. Yeasts encoding tumour antigens in cancer immunotherapy. Expert Opin Biol Ther 2005;5:565–75.

23. Gaggar A, Coeshott C, Apelian D et al. Safety, tolerability and immunogenicity of GS-4774, a hepatitis B virus-specific therapeutic vaccine, in healthy subjects: a randomized study. Vaccine 2014;32:4925–31.

24. Galazka A, Milstien J, Zaffran M. Thermostability of Vaccines.Global Programmefor Vaccines and Immunization (World Health Organization, Geneva) (1998).

25. Garmise RJ, Staats HF, Hickey AJ. Novel dry powder preparations of whole inactivated influenza virus for nasal vaccination. AAPS Pharm Sci Tech 2007;8:2–10.

26. Gleeson MA, White CE, Meininger DP et al. Generation of protease-deficient strains and their use in heterologous protein expression. Methods Mol Biol 1998;103:81–94.

27. Gould SJ, McCollum D, Spong AP et al. Development of the yeast *Pichia pastoris* as a model organism for a genetic and molecular analysis of peroxisome assembly. Yeast 1992;8:613–628.

28. Haller AA, Lauer GM, King TH et al. Whole recombinant yeast-based immunotherapy induces potent T cell responses targeting HCV NS3 and Core proteins. Vaccine 2007;25:1452–63.

29. Hassett KJ, et al. Glassy-State Stabilization of a Dominant Negative Inhibitor Anthrax Vaccine Containing Aluminum Hydroxide and Glycopyranoside Lipid A Adjuvants. J. Pharm. Sci. 2015;104:627–639.

30. Hassett KJK, et al. Stabilization of a recombinant ricin toxin A subunit vaccine through lyophilization. Eur. J. Pharm. Biopharm. 2013;85:279–86.

31. Hill A, Kilgore C, McGlynn M et al. Improving global vaccine accessibility. Current Opinion in Biotechnology 2016;42:67–73.

32. Klionsky DJ, Cuervo AM, Seglen PO. Methods for monitoring autophagy from yeast to human. Autophagy 2007;3:181–206.

33. Kumar R, Kumar P. Yeast-based vaccines: New perspective in vaccine development and application. FEMS Yeast Research 2019;19:1–22.

34. Kumar R, Rahman MA, Nazarko TY. Nitrogen starvation and stationary phase lipophagy have distinct molecular mechanisms. International Journal of Molecular Sciences 2020;21:1–13

35. Kumar R. Investigating the long-term stability of protein immunogen(s) for whole recombinant yeast-based vaccines. FEMS Yeast Research 2018;18:1–13.

36. Kumar R. Simplified protocol for faster transformation of (a large number of) *Pichia pastoris* strains. Yeast 2019;36:399–410

37. Kumru OS, Joshi SB, Smith DE et al. Vaccine instability in the cold chain: mechanisms, analysis and formulation strategies. Biologicals 2014;42:237–59

38. Kurtzman CP. Biotechnological strains of Komagataella (Pichia) pastoris are Komagataella phaffii as determined from multigene sequence analysis. Journal of Industrial Microbiology & Biotechnology 2009;36:1435–8.

39. Lee BY, Wedlock PT, Haidari LA et al. Economic impact of thermostable vaccines. Vaccine 2017;35:3135–3142.

40. Leung V, Mapletoft J, Zhang A et al. Thermal Stabilization of Viral Vaccines in Low-Cost Sugar Films. Sci Rep 2019;9:7631.

41. Liao YH, Brown MB, Martin GP. Investigation of the stabilisation of freeze-dried lysozyme and the physical properties of the formulations. Eur. J. Pharm. Biopharm. 2004;58:15–24.

42. Lloyd J, Lydon P, Ouhichi R et al. Reducing the loss of vaccines from accidental freezing in the cold chain: the experience of continuous temperature monitoring in Tunisia. Vaccine 2015;33:902–907.

43. Lovalenti PM, et al. Stabilization of live attenuated influenza vaccines by freeze drying, spray drying, and foam drying. Pharm. Res. 2016;33:1144–1160.

44. Lu Y, Bellgrau D, Dwyer-Nield LD et al.. Mutation-selective tumor remission with Ras-targeted, whole yeast-based immunotherapy. Cancer Res 2004;64:5084–8.

45. Maa YF, Ameri M, Shu C et al. Influenza vaccine powder formulation development: Spray-freeze-drying and stability evaluation. J. Pharm. Sci 2004;93:1912–1923.

46. Milistien JB, Stanley ML, Peter FW. Development of a More Thermostable Poliovirus Vaccine. The Journal of Infectious Diseases 1997;175(Suppl 1):S247–53

47. Mistilis MJ, et al. Long-term stability of influenza vaccine in a dissolving microneedle patch. Drug Deliv. Transl. Res. 2017;7:195–205.

48. Nguyen P, Botyanszki Z, Tay P et al. Programmable biofilm-based materials from engineered curli nanofibres. Nat Commun 2014;5:4945.

49. Norrby E, Uhnoo I, Brytting M et al. Polio närmar sig utrotning [Polio close to eradication]. Lakartidningen. 2017;114:EPDT.

50. Ohtake S, et al. Room temperature stabilization of oral, live attenuated Salmonella enterica serovar Typhi-vectored vaccines. Vaccine. 2011;29:2761–71.

51. Ohtake S, et al. Heat-stable measles vaccine produced by spray drying. Vaccine. 2010;28:1275–84.

52. Patterson R, Eley T, Browne C et al. Oral application of freeze-dried yeast particles expressing the PCV2b Cap protein on their surface induce protection to subsequent PCV2b challenge in vivo. Vaccine 2015;33:6199–6205.

53. Pelliccia M, Andreozzi P, Paulose J et al. Additives for vaccine storage to improve thermal stability of adenoviruses from hours to months. Nat Commun 2016;7:13520.

54. Ramchuran SO, Mateus B, Holst O et al.. The methylotrophic yeast Pichia pastoris as a host for the expression and production of thermostable xylanase from the bacterium Rhodothermus marinus. FEMS Yeast Res. 2005;5:839–50.

55. Schiller JT, Lowy DR. Raising expectations for subunit vaccine. J Infect Dis. 2015;211:1373–5.

56. Sears IB, O’Connor J, Rossanese OW et al. A versatile set of vectors for constitutive and regulated gene expression in Pichia pastoris. Yeast 1998;14:783–90.

57. Song JG, Lee SH, Han H. The stabilization of biopharmaceuticals: Current understanding and future perspectives. J. Pharm. Investig 2017;47:475–496.

58. Stubbs AC, Martin KS, Coeshott C et al. Whole recombinant yeast vaccine activates dendritic cells and elicits protective cell-mediated immunity. Nat Med 2001;7:625–9.

59. Takeshige K, Baba M, Tsuboi S et al. Autophagy in yeast demonstrated with proteinase-deficient mutants and conditions for its induction. J Cell Biol 1992;119:301–11.

60. Towbin H, Staehelin T, Gordon J. Electrophoretic transfer of proteins from polyacrylamide gels to nitrocellulose sheets: Procedure and some applications. PNAS USA 1979;76:4350–4354.

61. Vartak A, Sucheck SJ. Recent Advances in Subunit Vaccine Carriers. Vaccines (Basel). 2016;4:12.

62. Versteeg L, Almutairi MM, Hotez PJ et al. Enlisting the mRNA Vaccine Platform to Combat Parasitic Infections. Vaccines (Basel) 2019;7:122.

63. Wang G, Cao RY, Chen R et al. Rational design of thermostable vaccines by engineered peptide-induced virus self-biomineralization under physiological conditions. PNAS USA 2013;110:7619–7624.

64. Wang SH, Kirwan SM, Abraham SN et al Stable dry powder formulation for nasal delivery of anthrax vaccine. J. Pharm. Sci 2012;101:31–47.

65. Xiang SD, Scholzen A, Minigo G et al. Pathogen recognition and development of particulate vaccines: does size matter? Methods 2006;40:1–9.

66. Zhang W, Sack DA. Current Progress in Developing Subunit Vaccines against Enterotoxigenic Escherichia coli-Associated Diarrhea. Clin Vaccine Immunol 2015;22:983–91.

